# Sensory origin of visually evoked activity in auditory cortex: evidence for true cross-modal processing

**DOI:** 10.1101/2024.12.18.629217

**Authors:** Timothy Olsen, Andrea Hasenstaub

## Abstract

The meaning and functions of cross-modal sensory processing in the cortex is at the center of an ongoing debate. While some studies claim that such responses reflect genuine multisensory integration, others argue they are mere artifacts of stimulus-evoked movement or changes in internal state. We examined this issue by measuring face movements and neural activity in awake mouse primary auditory cortex (A1) and primary visual cortex (V1) during visual and auditory stimulation. Visual stimuli rarely evoked face movements, A1 responses to visual input remained robust even in the absence of movement, and optogenetic silencing of V1 reduced A1 visual responses, confirming a sensory origin of these cross-modal responses. These findings directly challenge the argument that cross-modal responses are purely movement-driven and emphasize that rather than assuming all cross-modal effects are artifactual, researchers must rigorously test each case.

**Highlights:**

- Unlike auditory stimuli, visual stimuli rarely evoke face movements
- Face movements explain sound-evoked firing in VC but not visually evoked firing in AC
- Silencing VC suppresses visually evoked firing in AC
- These results confirm a sensory (rather than motor) origin of these responses

## Introduction

For decades, sensory neuroscience was guided by the principle that primary sensory cortices are largely specialized for processing information from a single modality^1,2^, and that cross-modal convergence happens relatively late in the sensory processing hierarchy^3–6^.

Conversely, later studies reported cross-modal responses even in early sensory cortical regions^7–12^, creating an ongoing debate about the extent to which the earlier models of cortical sensory processing as hierarchical, segregated, and parallel accurately describe brain function. A recent and controversial contribution to the debate came from studies in awake mice which showed that intermittent sound stimuli frequently elicit face and whisker movements, leading to neural responses in visual cortex (VC) that can be misinterpreted as being evoked by the auditory stimulus itself rather than by the associated movement^13–15^.

These findings have been interpreted as supporting the idea that all or most cross-modal responses in sensory cortices should be reinterpreted as movement-related artifacts^15^, which itself supports the earlier idea that primary cortical sensory processing is fundamentally hierarchical and unimodal.

It is thus critical to revisit claims of cross-modal activity in other sensory regions, including the auditory cortex (AC). Previous studies^9,16–18^, including our own work^19–21^, have demonstrated that AC neurons can respond to visual stimuli. However, the extent to which these responses reflect genuine visual processing as opposed to movement artifacts has not been systematically tested. If visual responses in AC arise from stimulus-evoked movements, then they should be tightly linked to the presence of such movements. On the other hand, if these responses persist even in trials where no movement occurs, this would indicate a sensory-driven cross-modal response rather than an artifact.

To address this question, we used a combination of behavioral tracking, electrophysiological recordings, and optogenetic manipulations. First, we examined whether visual stimuli elicit face movements at rates comparable to auditory stimuli. Next, we assessed whether visually evoked activity in primary auditory cortex (A1) correlates with face movements, analyzing neuronal firing across trials with and without movement. Finally, we silenced the primary visual cortex (V1) optogenetically to determine whether visually driven responses in A1 depend on direct sensory input. We found that visual stimuli rarely induce face movements, and when they do, these movements do not explain visually evoked firing in A1. Instead, optogenetically silencing V1 suppresses visual responses in A1, consistent with a sensory rather than behavioral origin. These findings contradict the claim that face movements broadly explain cross-modal activity across sensory cortices, and show that visually evoked firing in A1 is genuinely sensory, not a movement artifact.

## Results

### Visual stimuli are much less likely to evoke face movements than auditory stimuli

We first tested whether similarly infrequent visual stimuli also tend to evoke face movements. Awake mice (Figure 1A) were, on randomly interleaved trials, presented with contrast-modulated noise movies (Figure 1B, red, 100 trials) or repeating noise burst stimuli (Figure 1B, blue, 30 trials), with each trial separated by 4-12 seconds. Face movements were quantified using Facemap software (Figure 1C)^22^, which applies singular value decomposition to capture face movements over time using multiple principal components (PCs). We initially focused on PC1, with further analysis of PCs 2-3 provided in the supplementary material to confirm our findings.

**Figure 1.**
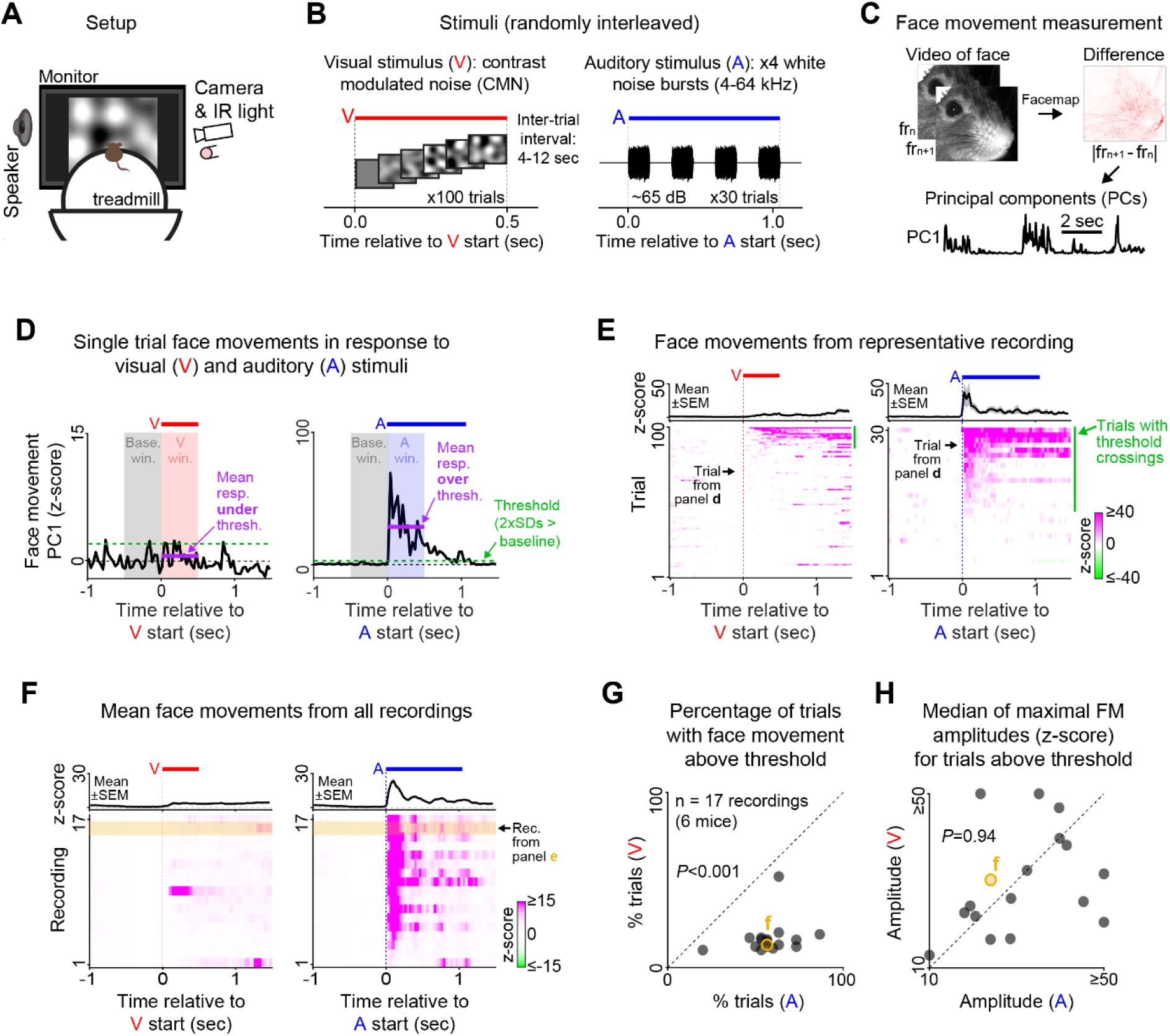
Visual stimuli are much less likely to evoke face movements than auditory stimuli. (**A**) Experimental setup. Mice were head-fixed and placed on a treadmill. Visual and auditory stimuli were presented while their face movements were recorded using an infrared camera and light. (**B**) Visual stimuli (V) were 0.5-second contrast modulated noise movies (left) and auditory stimuli (A) were 1.05-second repeating noise burst stimuli. (**C**) Face movements were measured using Facemap software, which calculated the absolute difference in pixel intensity between consecutive frames and extracted the first 500 principal components over time from this difference movie. (**D**) Face movements (PC1) in response to both stimulus types from representative trials. The red (left, V. win) and blue (right, A. win) shading shows the visual and auditory response windows, where visually- and sound- evoked face movements were measured. (**E**) Trial-aligned face movements (PC1) from representative recordings (bottom) and the mean of these face movements across trials (top). (**F**) Mean face movements for each recording (bottom), and the grand means across recordings (top). (**G**) Percentage of auditory versus visual trials with face movements (PC1) above threshold (n = 17 recordings, 6 mice). Auditory stimuli were significantly more likely than visual stimuli to evoke above-threshold face movements (*** p = 2.9 x10^-4^, two-sided Wilcoxon signed-rank test). (**H**) Median face movement amplitudes for above-threshold trials, compared between visual and auditory stimuli. There was no significant difference between stimulus types (p = 0.94, two-sided Wilcoxon signed-rank test).

Face movements were much less likely to occur during visual stimulus trials than during sound stimulus trials (Figures 1D-F). To quantify this, we calculated the percentage of auditory and visual trials with stimulus-induced face movements. A trial was considered to have stimulus-induced face movement if the mean face movement within 500 ms of stimulus onset exceeded 2 standard deviations from baseline (threshold shown by the green dashed line in Figure 1D; see Figure S1 for threshold rationale), where baseline is defined as the 500 ms prior to stimulus onset (gray shading in Figure 1D). The percentage of trials with face movement was substantially higher for auditory stimuli than for visual stimuli (Figure 1G, visual: 16.5 ± 9.6%, auditory: 58.0 ± 13.9%, two-sided Wilcoxon signed-rank test: *** p = 2.9 x 10-4). We also examined whether the face movements elicited by visual and auditory stimuli were of similar magnitudes, and found that they were (Figure 1H, visual: 31.0 ± 14.4, auditory: 32.4 ± 14.8, two-sided Wilcoxon signed-rank test: p = 0.94).

To ensure that our results were not dependent on our specific experiment and analysis choices, we ran further analyses using additional face movement principal components (Figure S2), and a different auditory/visual stimulus paradigm (Figure S3). In these complementary analyses, visual stimuli remained much less likely to evoke face movements than sound stimuli.

### Most visual responses in auditory cortex are not explained by face movements

Although visually evoked face movements were rare, they were not nonexistent, leading us to question how much these movements might explain visually-evoked activity in the auditory cortex. While recording face movements from head-fixed mice in response to visual stimuli (Figure 1A), we also recorded electrophysiological data from the primary auditory cortex (Figures 2A-C). Consistent with our previous studies^19,21^, about 8% of recorded cells in the auditory cortex responded with significantly elevated firing in response to visual stimuli (Figure 2D, red; n = 39 of 499 cells, p < 0.01, FDR corrected, two-sided Wilcoxon signed rank tests), most of which were located in the deep layers. When comparing visually evoked firing and face movements on the same trials (Figure 2E), in most AC cells, face movements appeared unrelated to visual responses (i). Yet for a smaller subset of cells, visually evoked firing did appear to accompany face movements (ii). To quantify the prevalence of such face motion artifacts and the effect of face movement on visually evoked firing, we compared the change in firing rate following the visual stimulus between trials with and without face movement, for all visually responsive cells (Figure 2F). As in Figure 1, we used a 2x SD threshold to detect trials with face movement. In only 20% of cells were visual responses significantly different between trials with and without face movement (Figure 2F, left, orange, 20%, p < 0.01, FDR corrected, two-sided Mann-Whitney U tests). In the vast majority (80%) of cells, the presence or absence of face movement made no difference to the visual responses (Figure 2F, left, gray). We further split these data by recording and mouse to determine if the effect of face movements on visually evoked firing depended on these factors (Figure 2F, right), but we found no such dependency; the visually responsive cells affected by face movements were randomly distributed across recordings (p = 0.21, Kruskal-Wallis test) and mice (p = 0.16, Kruskal-Wallis test).

**Figure 2.**
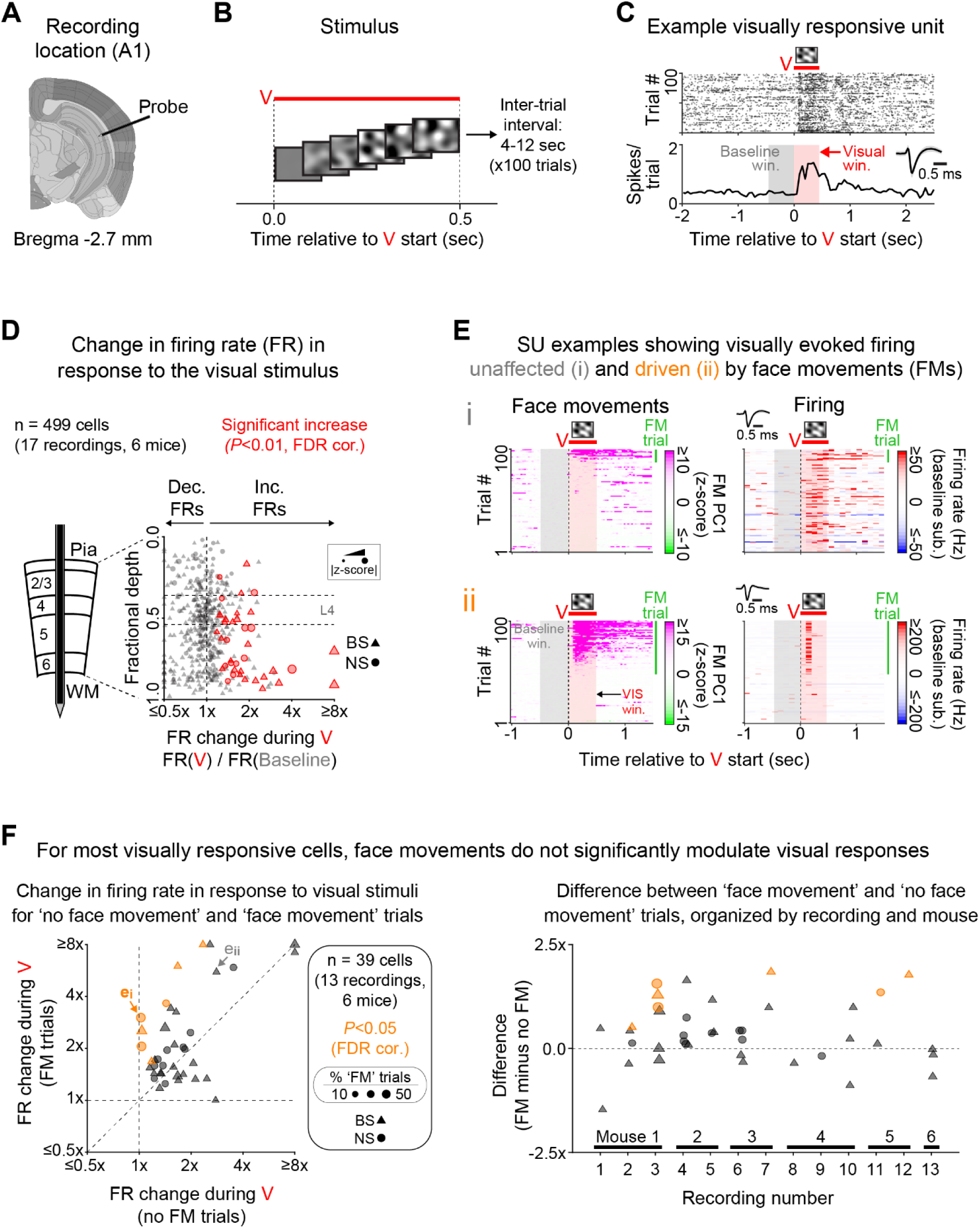
Most visual responses in auditory cortex are not explained by face movements. (**A**) Recording location. SU firing was recorded from all layers of the primary auditory cortex. (**B**) For these experiments, we used visual stimuli only. These consisted of 100 contrast-modulated noise movies, each lasting 500 ms and separated by 4-12 seconds. (**C**) Raster plot (top) and PSTH (bottom) showing significantly elevated firing in response to the visual stimulus for an example cell. Significance was determined using a two-sided Wilcoxon signed-rank test comparing single-trial spike counts in the baseline (gray) and visual (red) windows (p < 0.01, FDR corrected). (**D**) Change in firing rate in response to the visual stimulus plotted against cell depth for all recorded cells (n = 499 cells, 17 recordings, 6 mice). Cells with significant increases in firing are shown in red (p < 0.01, FDR corrected). (**E**) Face movements (left) and SU spike counts (right) for two example recordings/cells (i and ii). The top example (i) shows a cell with firing largely unrelated to face movements. The bottom example (ii) shows a cell with firing strongly related to face movements. The green bars to the right of each plot indicate trials where face movements (PC1) in the visual window (red shading) exceed the 2x SD threshold. Trials (rows) are ordered by maximal face movement amplitudes in the visual window. (**F**) (left) Change in firing rate for trials with (y-axis) and without (x-axis) face movement, for all cells with significant increases in firing in response to the visual stimulus (red cells in panel **D**). Cells showing a significant difference between the two trial types are marked in orange (p < 0.05, FDR corrected, two-sided Mann- Whitney *U* tests). Most cells did not show a significant difference between trials with and without face movements (gray data points, 32 out of 39: 82%). (right) Effects of face movements split by recording and mouse. The plot shows the difference in firing rate change for face movement minus no face movement trials (y- minus x-axis values in Figure 2F, left). We found no significant effect of recording (p = 0.21, Kruskal-Wallis tests) or mouse (p = 0.16, Kruskal-Wallis tests).

We confirmed that these results remained consistent when incorporating additional face movement principal components (Figure S4), and they did not differ when only including recording sites with onset latencies consistent with primary auditory fields (Figure S5). We also performed an additional analysis using a generalized linear model to compare the effects of visual stimuli and face movements on visual responses in the auditory cortex (Figure S6). This analysis showed that visual responses in the auditory cortex were more strongly related to visual stimuli than to face movements, further supporting our findings in Figure 2F.

In summary, most visual responses in the auditory cortex are not explained by face movements.

### Within the same experiment, face movements modulate cross-modal activity in the VC but not in the AC

Our findings indicate that stimulus-driven face movements do not explain cross-modal activity in the auditory cortex, while previous research^14,15^ indicated that face movements are major drivers of apparently cross-modal activity in the visual cortex. To verify that this difference is not due to differences between labs in experimental design or stimuli, we recorded simultaneously from the primary auditory and visual cortices and compared the effect of face movement on visually-evoked activity in the auditory cortex (Figure 3A) with that of sound-evoked activity in the visual cortex (Figure 3B), in interleaved trials during the same experiment. Plotting the difference in firing rate between trials with and without face movement revealed that visually-evoked firing in auditory cortex mostly remained stable across trials with and without face movements (Figure 3C, left), whereas sound-evoked firing in visual cortex consistently increased on face movement trials (Figure 3C, right).

**Figure 3.**
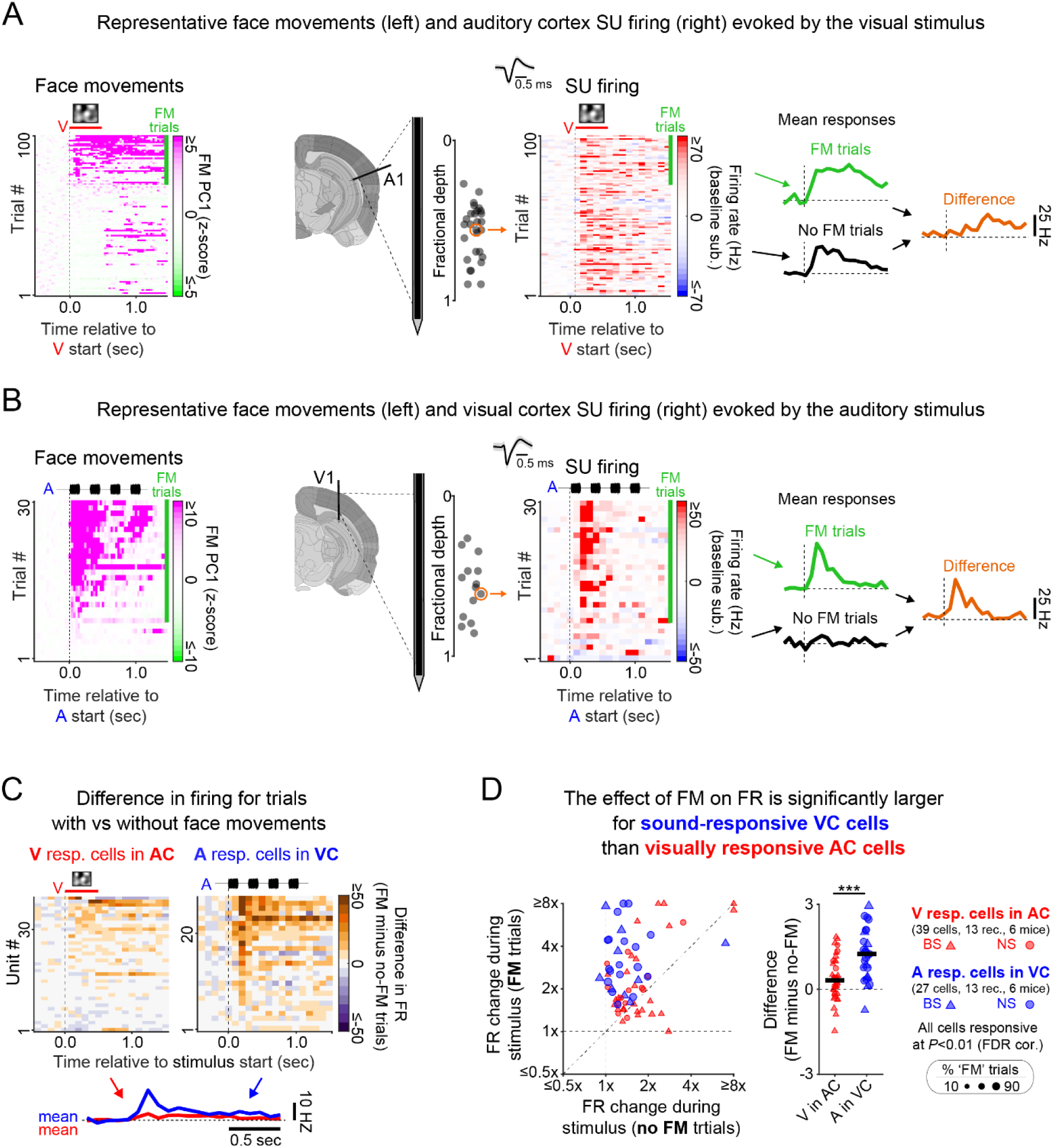
The effect of face movements on cross-modal activity differs between auditory and visual cortices. (**A**) Face movements (left) and SU firing (right) for a visually responsive unit in the auditory cortex. Mean firing differences between trials with and without face movement (far right, orange line) were minimal for this unit. (**B**) Same as **A**, but for a sound-responsive unit in the visual cortex, where face movement had a substantial effect on stimulus-evoked firing. (**C**) Firing rate differences between trials with and without face movement. Most visually responsive cells in the auditory cortex showed minimal differences (left), while sound-responsive cells in the visual cortex typically fired more during trials with face movement (right). (**D**) Firing rate changes in response to the stimuli, for visually responsive cells in the auditory cortex (red) and sound-responsive cells in the visual cortex (blue), plotted for trials with (y-axis) and without (x-axis) face movement (left). For most visually responsive cells in the auditory cortex, firing rate changes clustered near the diagonal, indicating a minimal effect of face movement. In contrast, firing rate changes landed substantially above the diagonal for nearly all sound-responsive cells in the visual cortex, indicating a positive, modulatory effect of face movements. Differences in firing rate changes (right) between face movement and no face movement trials were near zero for visually responsive cells in the auditory cortex, but significantly greater and predominantly positive for sound-responsive cells in the visual cortex (AC: 0.38 ± 0.74, VC: 1.2 ± 0.9, *** p = 1.6 x 10^-4^, two-sided Mann-Whitney *U* test).

To quantify this difference, we plotted the stimulus-evoked firing rate change for trials with (y-axis) and without (x-axis) face movement (Figure 3D, left), for auditory cortex cells with significant visually evoked firing (red) and visual cortex cells with significant sound-evoked firing (blue). This showed that while visually evoked firing in the auditory cortex typically remained stable across trials with and without face movements (red), sound-evoked firing in the visual cortex almost always increased during face movement trials. When comparing the difference in firing rate change between trials with and without face movement (‘with’ minus ‘without’), this difference was near zero for visually responsive cells in the auditory cortex (Figure 3D, right, red) but nearly always above zero for sound-responsive cells in the visual cortex (Figure 3D, right, blue), and this difference was statistically significant (AC: 0.4 ± 0.7, VC: 1.2 ± 0.9, two-sided Mann-Whitney *U* test: *** p = 1.6 x 10-4).

Taken together, these results show that while many apparently cross-modal responses in the visual system are explained by face movements, visually driven responses in the auditory system are generally not related to face movements.

### Sensory origin of visually-evoked activity in auditory cortex

As a final confirmation that visual activity in the AC is in fact driven by visual stimulation, we tested whether optogenetically silencing primary visual cortex (V1) could reduce it (Figures 4A-B). We selected V1 because essentially all visual information in the cortex comes by way of V1, and it is the gateway structure to other visual cortices^23^. In Ai32/PV-Cre mice (Figure 4A), we simultaneously recorded electrophysiological data from primary auditory and visual cortices, and used blue light to silence a large portion of primary visual cortex (Figure 4B). In visual cortex, blue light silenced nearly all broad-spiking cell firing (Figures 4C-D, brown), while narrow-spiking cell firing ranged from complete suppression to enhancement (Figure 4D, gray). Since most long-range cortical output comes from excitatory BS neurons^24^, this confirms that excitatory output from this area was effectively silenced. In auditory cortex, silencing visual cortex with blue light significantly suppressed visually evoked firing (Figures 4E-F, V only: 0.90 ± 0.80, V + opto: 0.40 ± 0.53, two-sided Wilcoxson signed-rank test: *** p = 5.4 x 10^-7^). We confirmed that this effect did not occur in mice lacking Ai32/PV-Cre expression (Figure S7), and was not due to regression to the mean (Figure S8). These results support a sensory rather than behavioral origin for visual activity in the auditory cortex.

**Figure 4.**
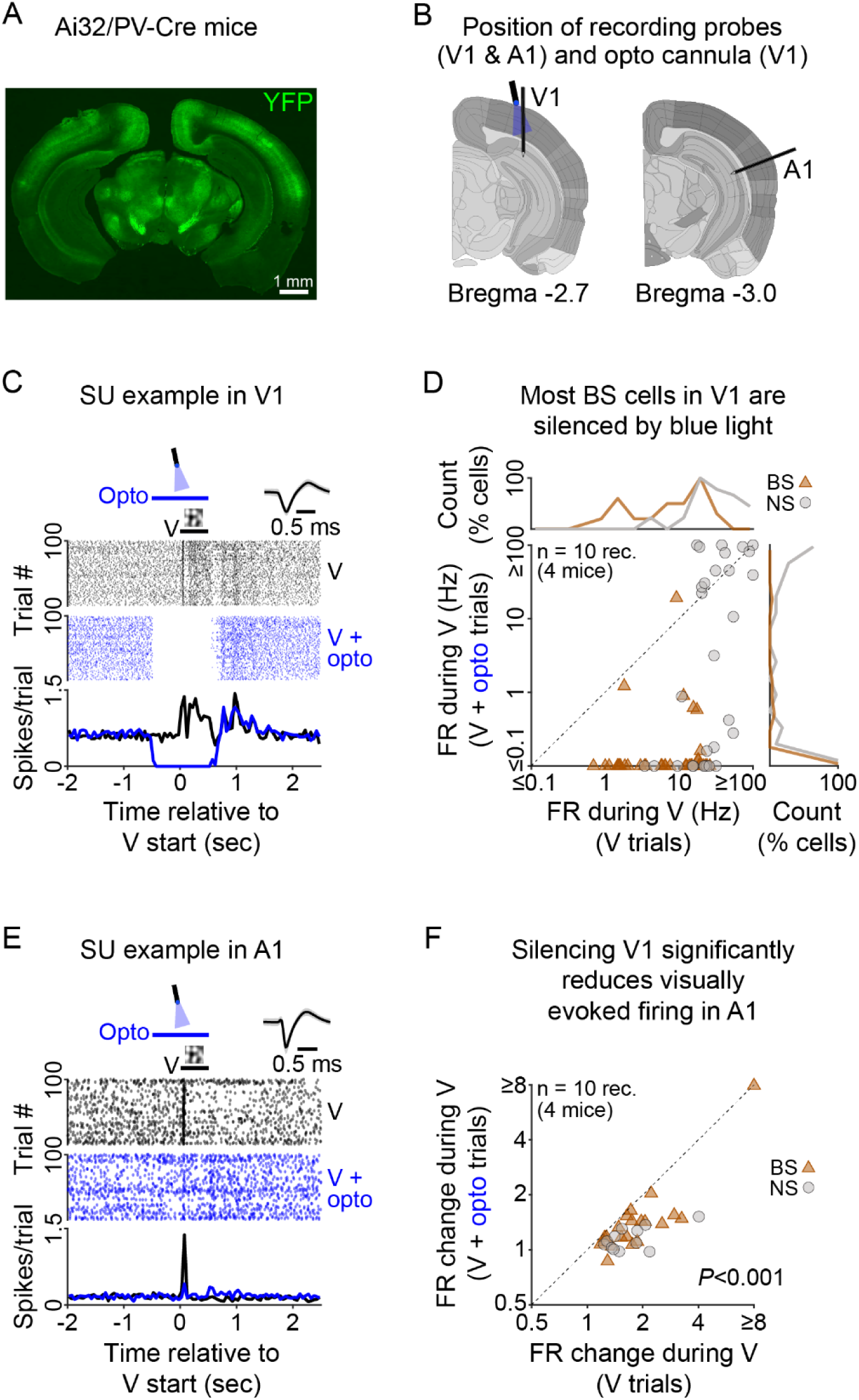
**Silencing visual cortex reduces visually evoked firing in auditory cortex**. (**A**) Histology showing eYFP fluorescence (green) from a representative Ai32/PV-Cre mouse (50 μm coronal slice). (**B**) Positioning of recording probes and opto cannula delivering blue light stimulation to the primary visual cortex. (**C**) Raster and PSTH showing firing from a representative BS cell in primary visual cortex, where its spontaneous and visually evoked firing was silenced by blue light stimulation. (**D**) Firing rate during the visual stimulus for trials with (y-axis) and without (x-axis) blue light stimulation, for all recorded BS (brown) and NS (gray) cells in primary visual cortex. (**E**) Raster and PSTH showing firing from a representative BS cell in primary auditory cortex, where visually evoked firing was suppressed following blue light stimulation of primary visual cortex. (**F**) Firing rate change in response to the visual stimulus for trials with (y-axis) and without (x-axis) blue light stimulation of primary visual cortex. Only cells with significantly elevated visually evoked firing in the control (no blue light) condition were included (same cells as in Figures 2F and 3D). Blue light stimulation significantly reduced visually evoked firing (V only: 0.90 ± 0.80, V + opto: 0.40 ± 0.53, two-sided Wilcoxson signed-rank test: *** p = 5.4 x 10^-7^).

## Discussion

Cross-modal activity has been reported in many sensory cortices^25–27^. Bimbard et al. (2023)^15^ showed that a significant portion of cross-modal auditory responses in the visual cortex were linked to movement, and hypothesized that similar effects might explain cross-modal responses throughout the cortex, including the visual responses in auditory cortex that we and many other groups have reported^9,16–19^. Our results challenge this interpretation. We found that standard visual stimuli rarely evoked face movements, whereas auditory stimuli did so much more frequently. Comparing trials with and without visually evoked face movements revealed that visually evoked activity in A1 was largely unaffected by face movement, with only 20% of visually responsive neurons showing significant firing rate differences between movement and non-movement trials. Furthermore, optogenetic silencing of V1 significantly reduced visually evoked activity in A1, confirming that these responses were sensory-driven rather than motor-related. While movement artifacts undoubtedly contribute to some cross-modal responses, our findings demonstrate that they cannot account for all such activity.

This suggests a fundamental difference in how cross-modal sensory information is processed in these two regions, with the visual cortex being more susceptible to movement artifacts and the auditory cortex exhibiting true visual sensitivity. Mechanistically, these differences may arise from variation in the effects of movement and arousal across primary sensory cortices^12^, which may be associated with variations in the projection patterns and post-synaptic effects of noradrenaline, acetylcholine, and other relevant neuromodulators^28^. Teleologically, these differences could reflect the sensory ecology of mice. Given their limited reliance on visual cues^29^, conventional visual stimuli may be less salient and thus less likely to trigger changes in arousal or body positioning than conventional sound stimuli. More salient visual stimuli, such as looming stimuli^30^, or quieter sounds that rarely evoke face movements^15^, could yield different outcomes. Nonetheless, these results highlight a distinct role for the auditory cortex in cross-modal sensory processing, and contribute to the growing body of research that challenges traditional sensory boundaries and suggest that the auditory cortex may be more visually responsive than previously appreciated.

The present findings are part of a broader wave of research over the past decades demonstrating cross-modal influences on sensory cortices, going beyond audiovisual interactions to demonstrate somatosensory effects in auditory cortex^11^, an influence of pain pathways in auditory cortex^31^, vestibular effects in visual cortex^10^, an influence of light on the olfactory bulb^32^, and so on. While our results confirm that some cross-modal effects are truly sensory-driven, they do not imply that all such effects are genuine. The key takeaway is that each case must be rigorously tested: apparent cross-modal activity may reflect real sensory integration, behavioral influences, or methodological artifacts, and distinguishing among these possibilities requires careful experimental design.

## Acknowledgements

We thank Professors Michael Stryker, Christoph Schreiner, and James Bigelow for helpful comments on this manuscript. This work was supported by the NIH grant R01NS116598, the Coleman Memorial Foundation, the PBBR Breakthrough Fund, and Hearing Research Incorporated. During the preparation of this work the authors used ChatGPT in order to improve paragraph and sentence flow. After using this tool/service, the authors reviewed and edited the content as needed and take full responsibility for the content of the publication.

## Methods

### Subjects

All procedures were approved by the Institutional Animal Care and Use Committee at the University of California, San Francisco. All mice were housed in cages of 2-5 under a 12 hr light/dark cycle, where food and water were provided *ad libitum*.

Experiments in all figures apart from Figure S3 were conducted on 6 mice (3 males, 3 females; mean age: 88 days, age range: 83-96 days). Of these, 4 were PV-Cre:Ai32 mice, which we obtained by crossing female PV-Cre mice (Jax strain: 012358, C57 background) with male Ai32 mice (Jax strain: 012569), and 2 were wild-type C57 mice (Jax strain: 000664; all on C57 background). For Figure S3, an additional set of 6 mice were used to validate our behavioral results with a different stimulus paradigm (3 males, 3 females; mean age: 98 days, age range: 74-106 days). This set included 4 PV-Cre:Ai32 mice and 2 SST-Cre mice (Jax strain: 013044, both on C57 background).

### Surgical procedures

For all mice, an initial surgery was performed under isoflurane anesthesia (4% for induction, 1-2% for maintenance) to fix a custom metal headbar to the skull with dental cement. Following this surgery, mice were allowed to recover for at least 2 days. Mice were then habituated to the recording setup for 20-90 mins at a time for 2-5 days. Mice then underwent a second surgery under isoflurane anesthesia where two ∼1-2 mm craniotomies were made. One was made to expose the primary auditory cortex (A1), and was made under the squamosal ridge at a location ∼2.2-3.2 mm posterior to bregma.

The other was made to expose the primary visual cortex (V1), and was centered at 3.0-3.5 mm posterior and 2.3-2.5 mm lateral to bregma (localization of the craniotomy above primary visual cortex was later verified stereotactically and based on responses to drifting gratings). Following this surgery, mice were allowed to recover overnight before electrophysiological recordings began. When not performing recordings, the craniotomies were covered with agarose (2%) and subsequently sealed with silicone elastomer (Kwik-cast, World Precision Instruments). Recordings were performed daily for up to 3 days following the craniotomies.

### Recording setup

All recordings were performed inside a sound attenuation chamber (Industrial acoustics), from awake mice placed on a spherical treadmill^33^. Visual stimuli were presented to the mice using a 19 inch monitor (ASUS VW199T-P or Dell P2016t, 60 frames per second, 1440 by 900 pixels) placed ∼25 cm directly in front of them, while auditory stimuli were delivered from a free-field electrostatic speaker (ES1, TDT) at 192 kHz. The speaker was placed immediately to the left of the monitor (relative to the mice), also ∼25 cm from the mice. To record face movements, we used a Mako U-130B camera (Allied Vision) with a 13-130 mm varifocal lens (B&H Photo Video), infrared light for illumination (Mightex SLS-0208-A), and custom software written in Matlab. The camera and infrared light were placed to the immediate right of the monitor (relative to the mice), and face videos were recorded at 30 frames per second.

### Electrophysiology

We recorded neural activity from the brain of awake mice using Neuropixels 1.0 probes. Probes were implanted into the primary auditory and visual cortices. The probes were continuously lowered into the brain at a speed of ∼1 μm/sec until silent channels were observed both above (pia) and below (white matter) activity in the cortex – typically to a depth of ∼1400 μm. Once this depth was reached, the recording sites were covered with agarose (2%) and the probes were left to settle for ≥30 minutes before recordings started. Data was recorded at a sampling rate of 30 kHz using a Neuropixels PXIe acquisition module (PXIE_1000), National instruments chassis (NI PXIe-1071), and SpikeGLX software (v.20230411). Single unit activity was obtained using Kilosort 2.0^34^ and results were curated using Phy (https://github.com/cortex-lab/phy). Splitting single units into broad and narrow spiking types, and determining their cortical depths, was performed using methods identical to those in our previous publication^35^. All spiking post-stimulus time histograms (PSTHs) were generated using 50 ms bins.

### Optogenetic silencing

We silenced a large portion of the primary visual cortex by shining blue light on this area in Ai32/PV-Cre mice. An optical fiber (Thor Labs, 400 μm tip diameter, 0.39 NA) was positioned just above the brain surface and parallel to the recording electrode in the primary visual cortex. This fiber was connected to a fiber-coupled LED light source (Mightex) and light was delivered at an intensity of 40 mW/mm^2^.

### Visual stimuli

Visual stimuli consisted of contrast modulated noise (CMN) movies, which are broadband stimuli designed to maximally activate the visual cortex^36^. The CMN was generated at 60x60 pixels and then interpolated to 900x900 pixels, covering an area of approximately 54 x 54° in the field of view. CMN stimuli were ramped from zero contrast (gray screen) to full contrast over 250 ms. The monitor during the inter-trial interval was black, thus giving an abrupt stimulus onset (black to gray). The monitor brightness was calibrated using a SpectraScan PR-670 device to 31 - 44 cd/m2 at gray screen.

### Sound stimuli

For most experiments, sound stimuli consisted of a repeated noise burst stimulus, which consisted of four 150 ms noise bursts, each separated by 150 ms of silence. The noise bursts were white noise band-pass filtered between 4 and 64 kHz, and were ramped on and off using 5 ms linear ramps. The same repeating noise burst stimulus was used for all trials. In some experiments (Figure S3), we used a random double sweep stimulus, which is a continuous, spectrally sparse sound that effectively drives neurons with diverse tuning properties in the primary auditory cortex^37^. For these experiments, a different random double sweep stimulus was presented on each trial. Both sound stimuli were presented at ∼65 dB, calibrated using a Brüel & Kjær 2209 and model 4939 microphone.

### Stimulus timing

All experiments, except for those in Figure S3, included three trial types: (1) visual stimuli (V; 100 trials), (2) visual stimuli with optogenetic stimulation (V + opto; 100 trials), and (3) auditory stimuli (A; 30 trials). Trials were randomly interleaved with inter-trial intervals of 4–12 seconds. In the V + opto trials, blue light stimulation started 0.5 seconds before the visual stimulus onset and ended with its offset. The experiment in Figure S3 included a single trial type with both visual and auditory stimuli. The auditory stimulus lasted 1 second, while the visual stimulus began 0.5 seconds before the start of the auditory stimulus and ended 0.3 seconds after it finished.

### Identifying recording sites with firing properties consistent with primary auditory fields

We only analyzed recordings showing clear signs of frequency tuning. Multi-unit (MU) firing was measured in response to 100 ms tones, using methods from our previous publication^35^, across 14 logarithmically spaced frequencies (4–64 kHz) and 7 attenuation levels (35–65 dB in 5 dB steps). For each recording channel, we calculated the frequency tuning curve (FTC), collapsed across attenuation, and measured its variance. This variance was compared to a null distribution of variance values generated by shuffling frequency labels (1000 iterations) to determine if it exceeded chance. A recording was classified as frequency-tuned if at least one-third of tone-responsive channels showed significant FTC variance (p < 0.01). Tone-responsive channels were identified by significantly elevated tone-evoked firing (collapsed across frequency and attenuation) compared to baseline, using a two-sided Wilcoxon signed-rank test (p < 0.01) on single-trial spike counts 100 ms before and during the tones.

We also assessed whether recordings had onset latencies consistent with primary auditory fields, which includes our target area (A1). These latencies are typically shorter than those in secondary auditory fields (5-18 ms vs. 8-32 ms^38^). For each channel, we measured the onset latency of MU responses to tones (collapsed across frequency and attenuation) by identifying when the response first exceeded 2.5 SDs of baseline firing (baseline: 100 ms prior to tone onset). As in our previous reports^21,35^, a recording was considered to be within a primary auditory field if the median onset latency across channels with significantly elevated tone-evoked firing was ≤14 ms. Since 82% of our recordings met this criterion, our results predominantly reflect firing from units in primary auditory fields. However, we also verified that our results remained unchanged when restricting analyses to these recordings (Figure S5).

### Face movement analysis

We used Facemap software to process face videos^22^, which applies singular value decomposition to quantify facial movement over time using multiple principal components (PCs). We used PC1 for most of our analyses, and then confirmed our results using additional principal components (PCs 2 & 3, Figures S2 & S4). For all analyses and plots, face movements were first aligned to visual and/or auditory stimuli for each trial. We then separately z-scored each trial by subtracting the mean baseline movement and dividing by the baseline standard deviation, using the baseline periods shown in each figure by gray shading. Trials were classified as face movement trials if mean face movement during the stimulus period (auditory or visual) exceeded 2 SDs (see Figure S1 for threshold rationale). All face movement plots have a time resolution of 33.3 ms, determined by the video frame capture rate of 30 frames per second.

### Generalized linear model

To confirm our main results, we used a generalized linear model (GLM) to assess how face movements contribute to visually evoked firing in the auditory cortex. The GLM attempted to predict spike counts within 500 ms windows, either during visual stimuli or during the inter-trial / baseline periods, using two predictor variables: a binary variable (Vstim) indicating whether spike counts came from visual (1) or baseline (0) windows, and the mean face movement (PC1) during each window (FM). The model equation is defined as follows, where ’win’ represents a 500 ms window:

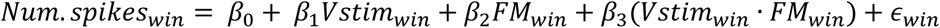

We ran a GLM for each recorded cell using Matlab’s (v.2024a) function fitglm, specifying ‘Poisson’ for the ‘distribution’ argument and ‘Log’ for the ‘Link’ argument.

### Statistics

All statistical analyses were performed in Matlab (v.2024a). We used a two-sided Wilcoxon signed-rank test for comparing two groups of paired data, and a two-sided Mann-Whitney *U* test for comparing two groups of unpaired data. For analyses involving multiple comparisons using these tests, such as across multiple cells (e.g., Figure 2D), we adjusted p-values using the Benjamini-Hochberg procedure to control for the false discovery rate (FDR). When comparing across more than two groups of unpaired data, we used a Kruskal-Wallis test.

---

**Figure S1.**
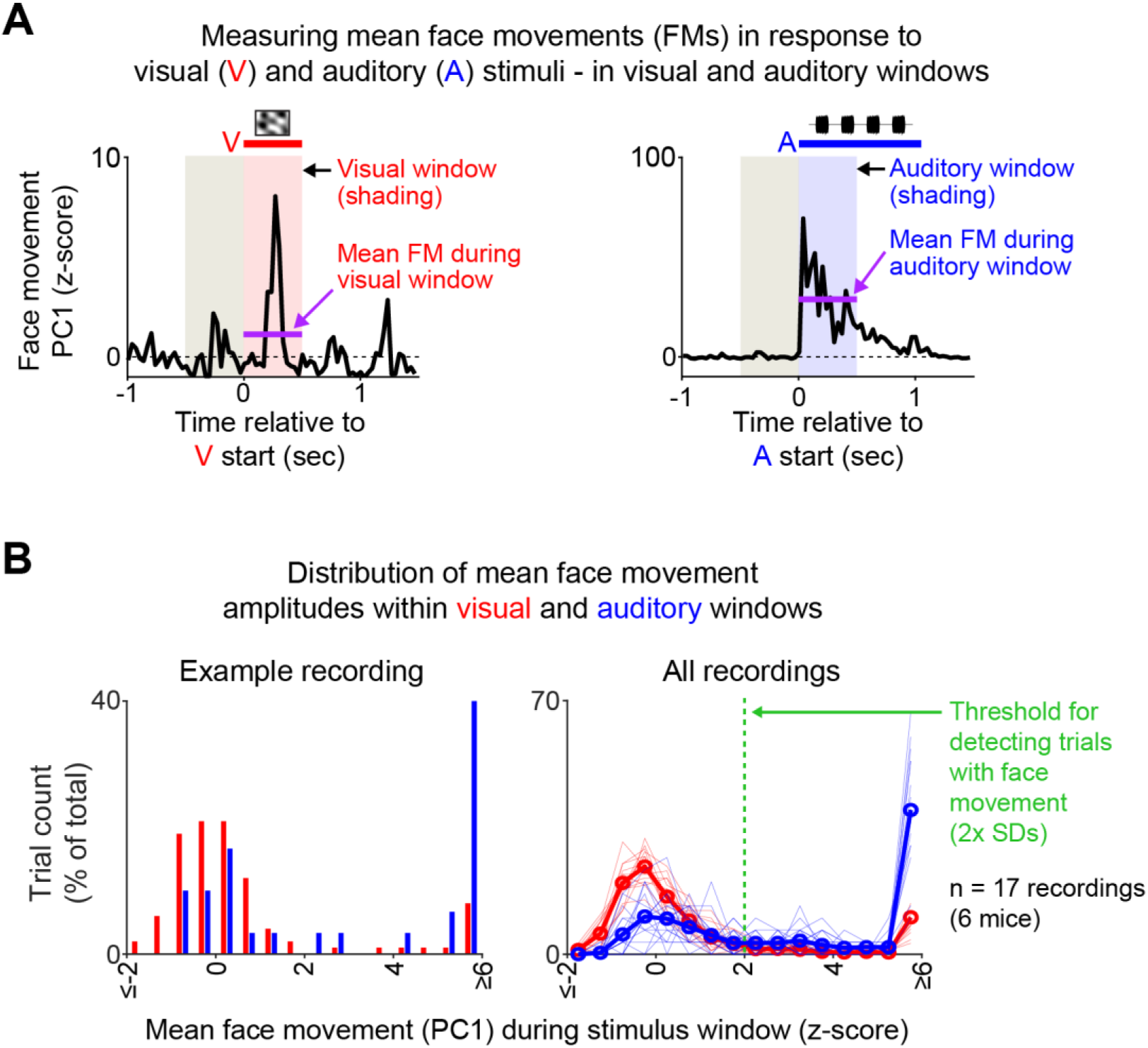
**Threshold selection for detecting trials with face movement**. (**A**) Single-trial face movements (PC1) from an example behavioral session showing mean stimulus-evoked responses (purple lines) during visual (red shading) and auditory (blue shading) trials. (**B**) Histograms showing the percentage of trials with various mean response amplitudes, for an example behavioral session (left) and for all sessions (right; thin lines, histograms for individual sessions; thick line, grand mean across sessions). Nearly all sessions showed a bimodal distribution of face movement magnitudes, with one peak near zero (presumably corresponding to noise) and another peak corresponding to stimulus-evoked face movements. Based on these, we chose a threshold of +2 SD for separating trials with and without stimulus-evoked face movements (green dashed line).

**Figure S2.**
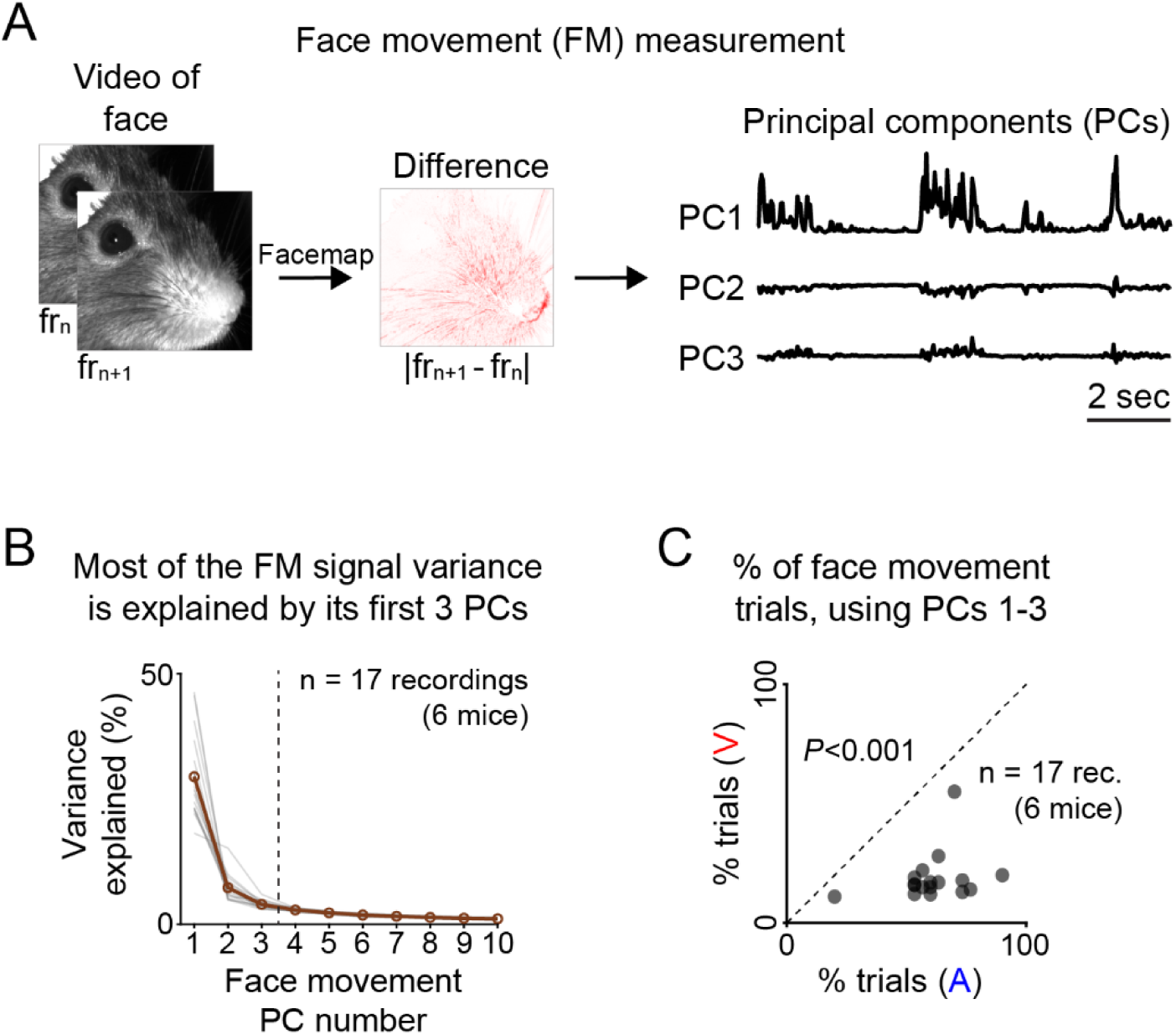
**Even with more face movement PCs analyzed, visual stimuli remain less likely to evoke face movements than auditory stimuli**. (**A**) Multiple face movement principal components (PCs) were obtained from face videos using Facemap software. (**B**) The percentage of variance explained by each face movement PC (up to 10), for individual behavioral sessions (thin gray lines) and their grand mean (thicker brown line with scatter points). PC1 consistently explained most of the variance, followed by far smaller contributions from PCs 2 and 3, and negligible contributions for PCs 4 and above. Following this observation, we tested if our results changed using PCs 1-3 rather than PC1 only (Figure 1). (**C**) Percentage of trials with above-threshold face movement (for at least one of PCs 1-3) for visual versus auditory stimuli. Even with additional PCs analyzed, the percentage of trials with above-threshold face movements remained significantly higher for auditory than for visual stimuli (V: 18.8 ± 10.2%, A: 61.0 ± 14.7%, two-sided Wilcoxon signed-rank test: *** p = 2.9 × 10⁻⁴).

**Figure S3.**
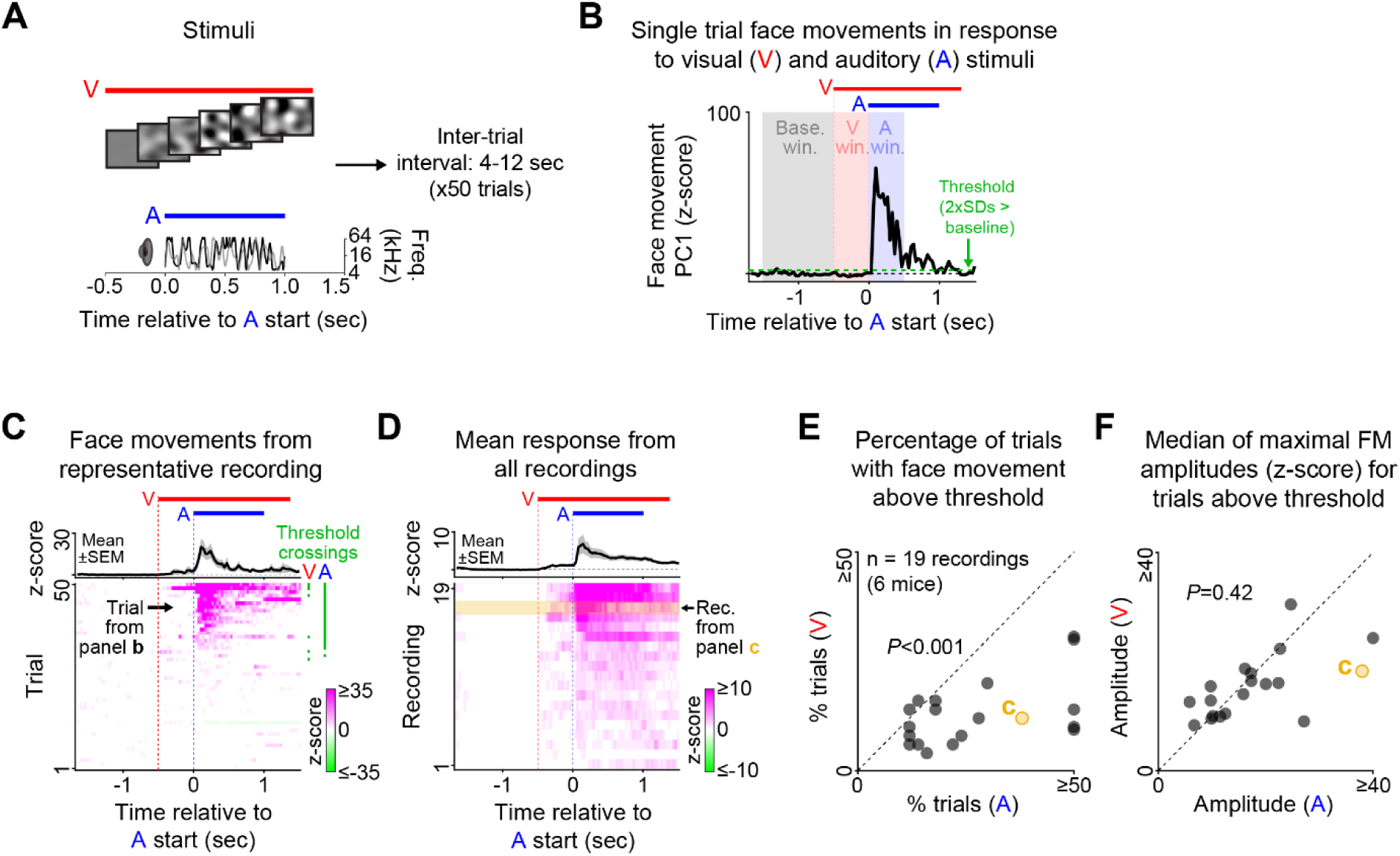
**Visual stimuli remain less likely than auditory stimuli to evoke face movements in a different stimulus paradigm**. We tested whether our main results from Figure 1 held true when using a different stimulus paradigm. (**A**) Stimulus trials included both visual (V) and auditory (A) stimuli. The visual stimuli were 50 different contrast-modulated noise movies, each lasting 1.75 seconds. The auditory stimuli were 50 different random double sweep sounds, each starting 0.5 seconds after the visual stimulus onset and lasting 1.0 second. Trials were separated by intervals of 4–12 seconds. (**B**) Face movement (PC1) in response to the visual + auditory stimulus from a representative trial. Shaded areas indicate the baseline window used to z-score the face movement traces (gray; separate z-scoring for each trial), and the visual (red) and auditory (blue) windows in which visual- and sound-evoked face movements were measured. (**C**) Trial-aligned face movements from a representative behavioral session, ordered by the maximal face movement. Green bars indicate trials on which movements exceeded the 2x SD threshold during the visual (V) or auditory (A) windows. The plot at the top shows the mean response across trials. (**D**) Mean face movements for all behavioral sessions (bottom), and their grand mean (top). Rows are ordered by maximal face movement. (**E**) Percentage of trials with face movements in the visual versus auditory windows. Similar to our previous stimulus paradigm (Figure 1G), this percentage was significantly higher for auditory than for visual stimuli (V: 12.9 ± 7.3%, A: 30.8 ± 21.6%, *** p = 2.8 × 10⁻⁴, two-sided Wilcoxon signed-rank test). (**F**) Median face movement amplitudes for above-threshold trials, compared between visual and auditory windows. Results are similar to our previous stimulus paradigm (Figure 1H): although visual and auditory stimuli differ in their propensity to evoke face movements, there is no significant difference between the magnitude of face movements evoked during the auditory and visual response windows (V: 15.4 ± 5.8, A: 18.4 ± 11.5, p = 0.42, two-sided Wilcoxon signed-rank test).

**Figure S4.**
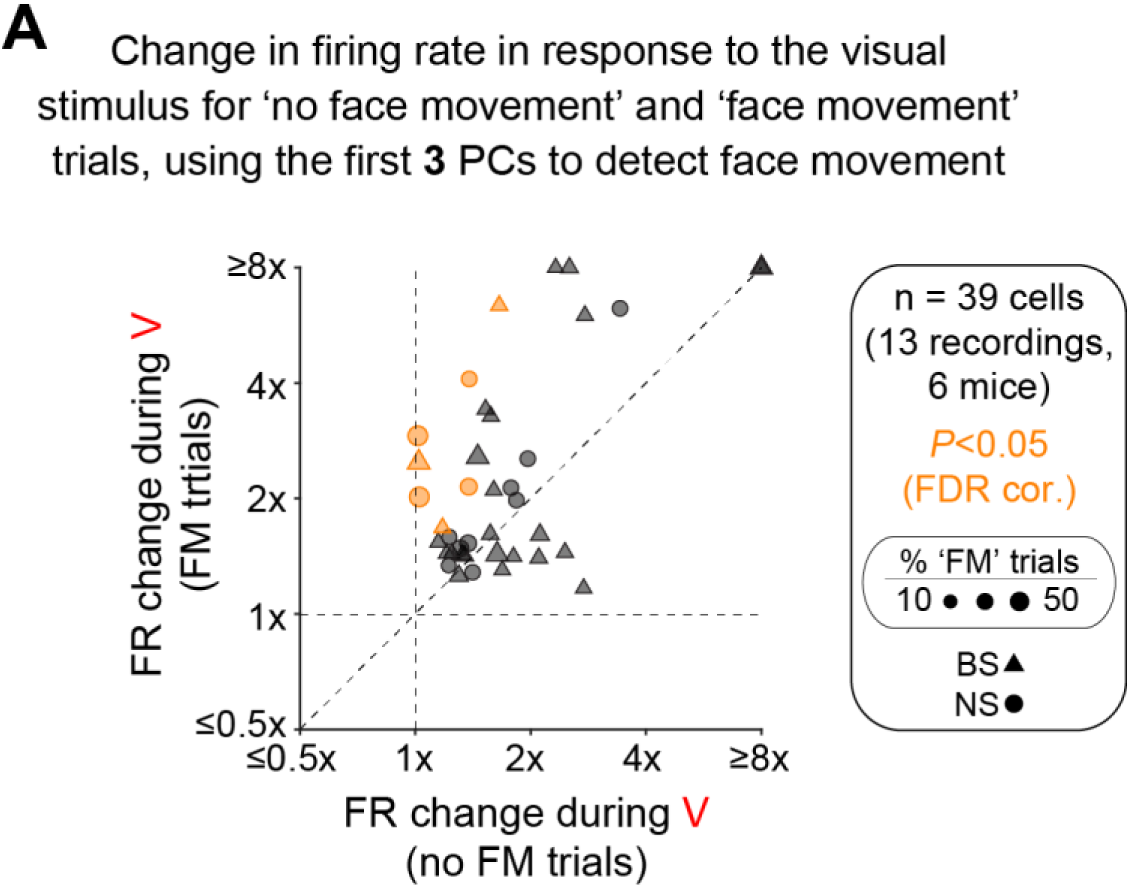
**Even when face movements are analyzed using more principal components, they still do not account for most visual responses**. (**A**) As in Figure 2F (left), but now face movement trials are defined based on movement in any of PCs 1-3. Consistent with our main results, most visually responsive cells did not show a difference in visually evoked firing between trials with and without face movement (gray data points, 31 out of 39, 79%).

**Figure S5.**
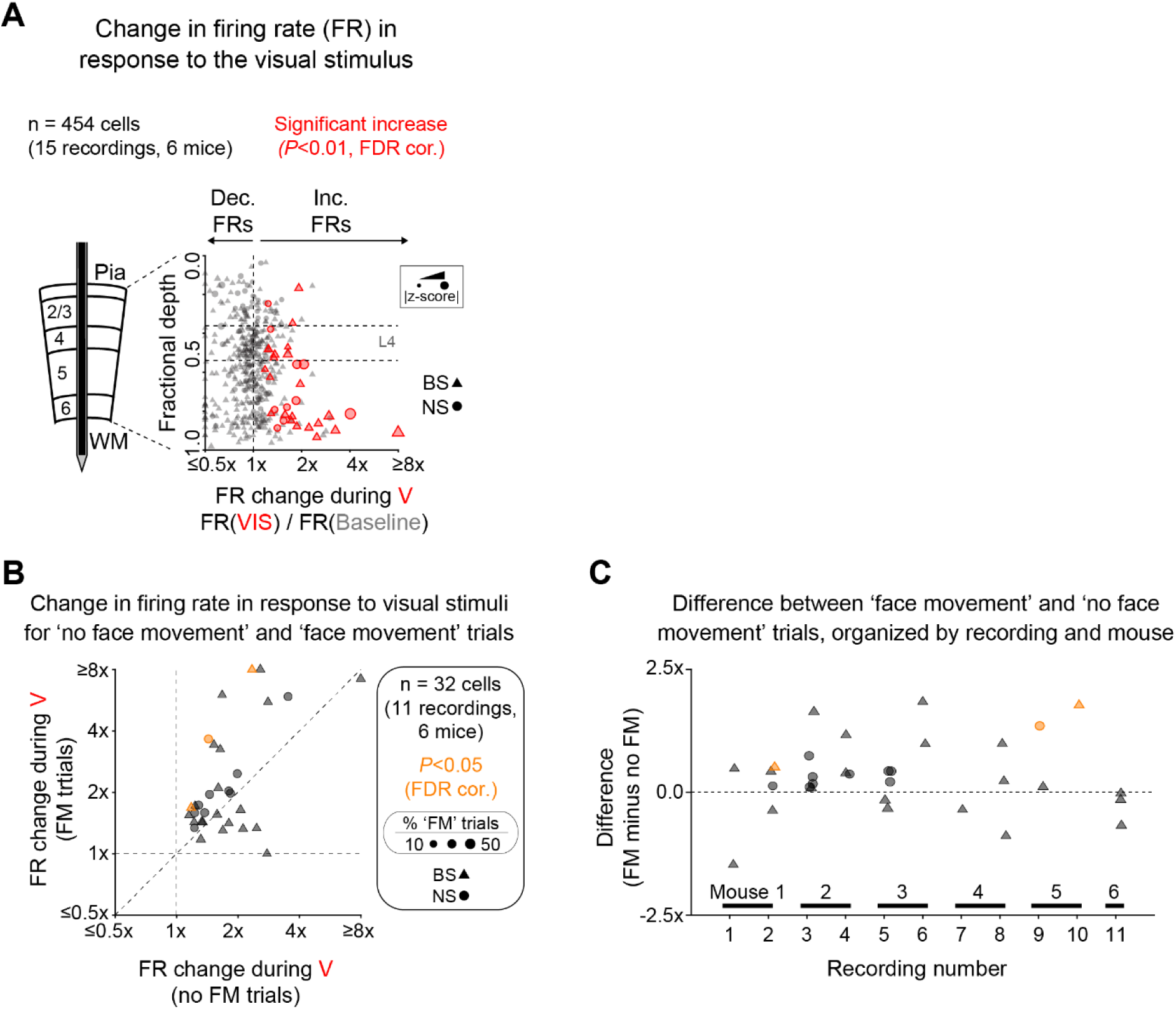
**Visual responses in primary auditory fields are largely independent of face movements**. To assess whether our results changed when only including units from recordings with firing properties consistent with primary auditory fields (see Materials and Methods for criteria), we reanalyzed our main findings using data from these recording sites only. In total, 15 out of 17 recordings (from 6 mice) had firing properties consistent with primary auditory fields. (**A**) From a total of 454 cells from these recordings (down from 499), 32 of these cells showed significantly elevated firing in response to the visual stimulus (down from 39 in Figure 2d), and these were still mostly found in the deep layers. (**B**) Out of these 32 visually responsive cells, 3 showed visually evoked firing that differed significantly between trials with and without face movements (orange data points; 9%, compared to 18% in Figure 2F, left). **c**, In addition, similar to Figure 2F (right), the effects of face movement on visually evoked firing were not dependent on recording (p = 0.22, Kruskal-Wallis test) or mouse (p = 0.20, Kruskal-Wallis test). These findings show that face movements still had little effect on visually evoked firing when only considering units from primary auditory fields.

**Figure S6.**
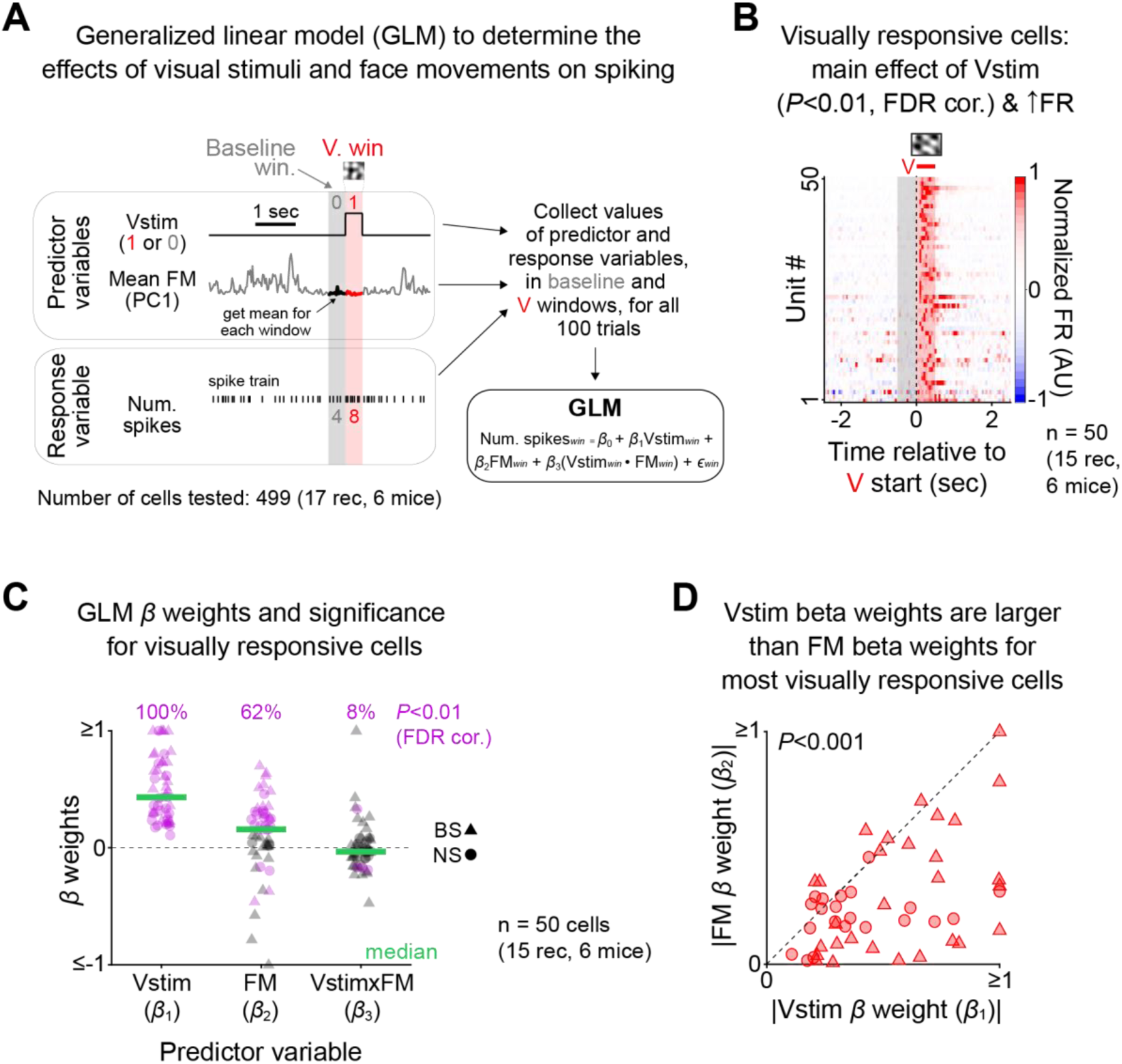
**A generalized linear model shows that visual responses are more related to the visual stimulus than to face movements**. To validate our main findings in Figure 2F using a complementary analysis we used a generalized linear model to analyze our data. (**A**) Generalized linear model design for each unit. The model attempted to predict the number of spikes during 500 ms windows (win), either during the visual stimulus (red shading) or during the preceding baseline periods (gray shading), based on two predictor variables: a binary variable (Vstim) indicating the window type, with 0 for baseline windows and 1 for visual windows, and the mean face movement (FM) in each window (PC1). The GLM equation is shown in the bottom right. We tested for main effects of Vstim and FM, and for an interaction between these two variables. (**B**) PSTHs showing firing in response to the visual stimulus for all visually responsive units, as determined by a significant main effect of Vstim (p < 0.01, FDR corrected), and an increase in firing relative to baseline (n = 50 cells, 15 recordings, 6 mice). Firing rates are normalized by their absolute maximum in the visual response window (red shading). (**C**) Model beta weights for the two predictor variables (Vstim and FM), and their interaction variable (Vstim x FM), are shown for all visually responsive cells (cells in panel **B**). The percentage of cells with significant beta weights (purple data points; p < 0.01, FDR corrected) was highest for Vstim (100%), lower for FM (62%), and minimal for Vstim x FM (8%). (**D**) Beta weights for the predictor variables Vstim and FM (absolute values) were compared on a cell-by-cell basis for all visually responsive cells. Vstim beta weights were significantly larger than FM beta weights (*** p = 4.7 × 10-7, two-sided Wilcoxon signed-rank test). These results, along with those in panel **C**, further indicate that face movements do not account for most visually evoked firing.

**Figure S7.**
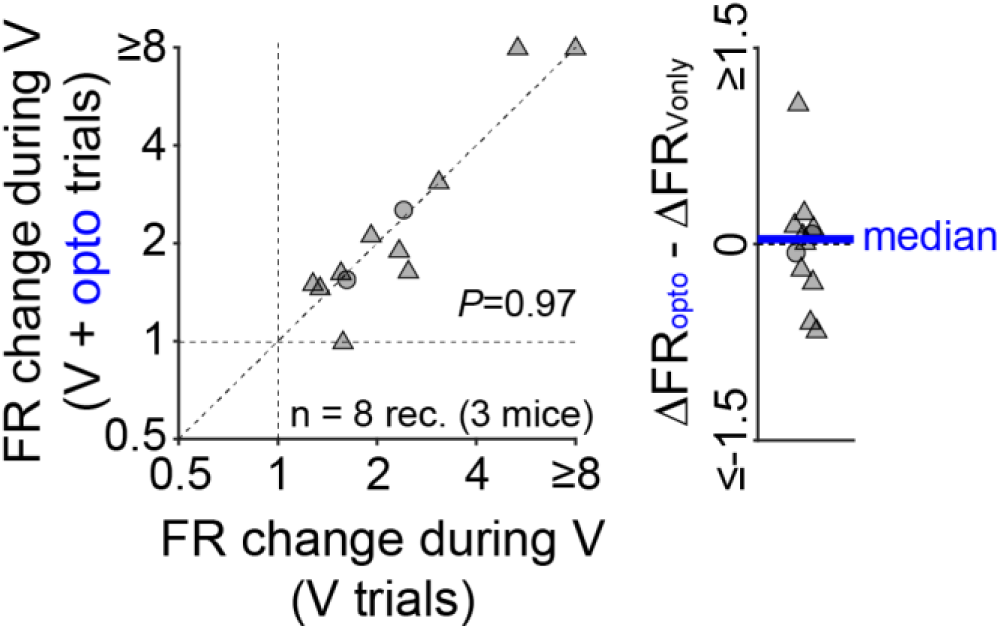
**Blue light stimulation of V1 does not affect visually evoked firing in A1 of Ai32/PV-Cre-negative mice**. Same as Figure 4F, but now in 3 mice lacking Ai32/PV-Cre expression. Two mice were wild-type C57 mice, and one was an Ai32/PV-Cre littermate lacking Ai32/PV-Cre expression. Blue light stimulation of V1 had no effect of visually evoked firing in A1 (V only: 1.24 ± 0.87, V + opto: 1.23 ± 1.06, two-sided Wilcoxon signed-rank test: p = 0.97). The plot on the right shows the y-axis values minus the x-axis values for the plot on the left.

**Figure S8.**
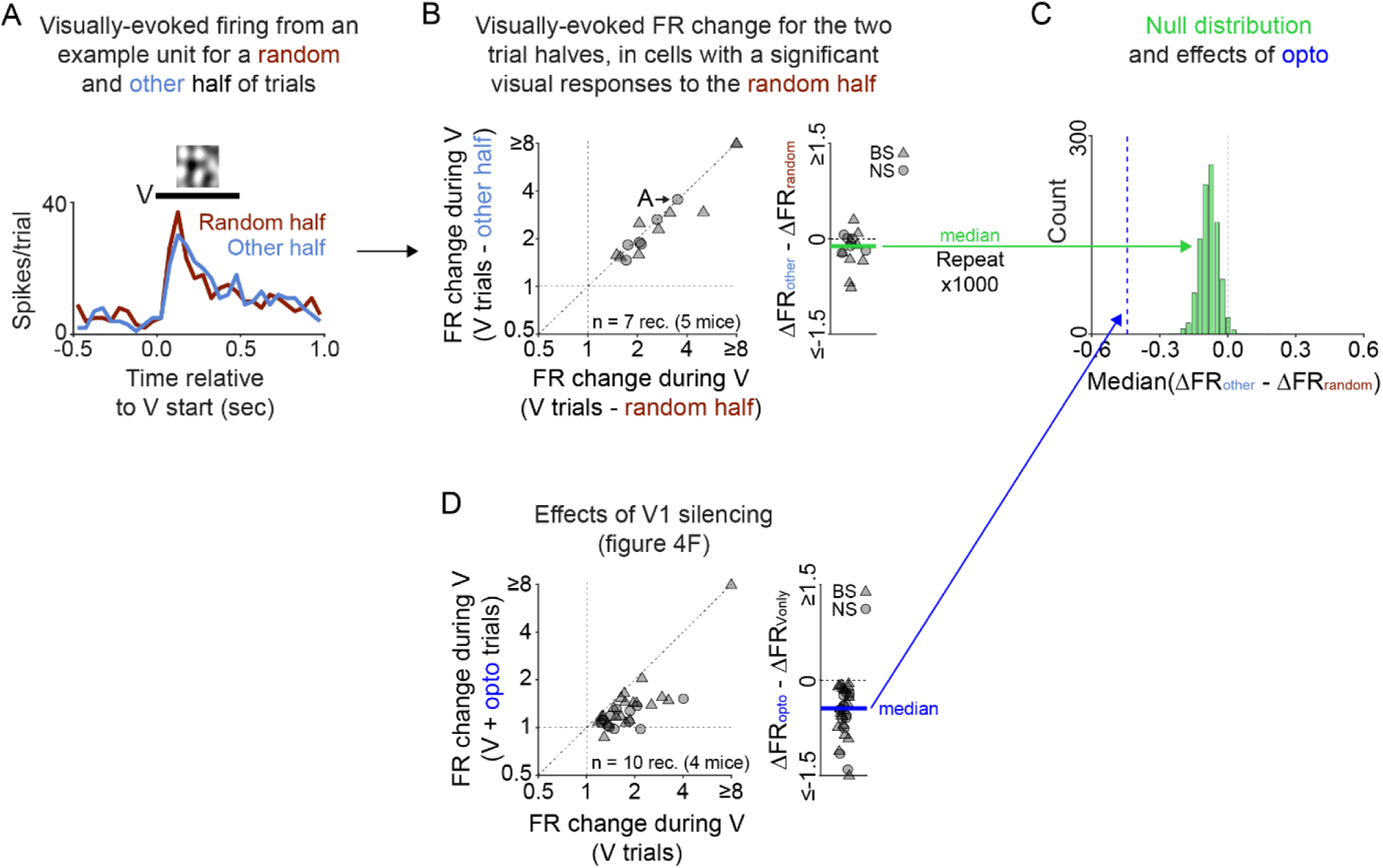
**The effect of V1 silencing on visual firing in A1 is not due to regression to the mean**. In Figure 4F, we observed a significant reduction in visually evoked firing in the auditory cortex following optogenetic silencing of V1. For that analysis, we included only cells with a significant increase in visually evoked firing in the visual-stimulus-only condition, which by itself would predict lower firing during the visual-stimulus-plus-opto condition due to regression to the mean. To confirm that the effects of V1 silencing were not solely explained by regression to the mean, we performed a control analysis. (**A**) For all recorded cells, we measured firing in response to a randomly chosen half of the visual-stimulus-only trials (50), and then separately for the other half of these trials (50). (**B-D**) Then, among cells showing a significant increase in visually evoked firing in the random half condition (p < 0.01, FDR-corrected, two-sided Wilcoxon signed-rank tests), we compared the firing rate change between the random and other halves (**B**, left). If the results in Figure 4F were solely due to regression to the mean, we would expect a similar magnitude decrease in visually evoked firing for the other compared to random half. To assess this, we took the differences in FR change for each cell (**B**, right, other minus random), computed the median of these differences (green bar), and repeated this procedure 1000 times to form a null distribution (**C**, green). Although a small decrease in FR change did occur in most iterations for the other vs random half of trials, it was much smaller than the decrease in FR change following V1 silencing (panel **D**, and blue line in **C**).

